# Multi-Molecular Hyperspectral PRM-SRS Imaging

**DOI:** 10.1101/2022.07.25.501472

**Authors:** Wenxu Zhang, Yajuan Li, Anthony A. Fung, Zhi Li, Hongje Jang, Honghao Zha, Xiaoping Chen, Fangyuan Gao, Jane Y. Wu, Huaxin Sheng, Junjie Yao, Dorota Skowronska-Krawczyk, Sanjay Jain, Lingyan Shi

**Author notes:** These authors contribute equally (Wenxu Zhang, Yajuan Li, Anthony A. Fung).

## Abstract

Lipids play crucial roles in many biological processes under physiological and pathological conditions. Mapping spatial distribution and examining metabolic dynamics of different lipids in cells and tissues in situ are critical for understanding aging and diseases. Commonly used imaging methods, including mass spectrometry-based technologies or labeled imaging techniques, tend to disrupt the native environment of cells/tissues and have limited spatial or spectral resolution, while traditional optical imaging techniques still lack the capacity to distinguish chemical differences between lipid subtypes. To overcome these limitations, we developed a new hyperspectral imaging platform that integrates a Penalized Reference Matching algorithm with Stimulated Raman Scattering (PRM-SRS) microscopy. With this new approach, we directly visualized and identified multiple lipid species in cells and tissues in situ with high chemical specificity and subcellular resolution. High density lipoprotein (HDL) particles containing non-esterified cholesterol was observed in the kidney, indicating that these pools of cholesterol are ectopic deposits, or have yet to be enriched. We detected a higher Cholesterol to phosphatidylethanolamine (PE) ratio inside the granule cells of hippocampal samples in old mice, suggesting altered membrane lipid synthesis and metabolism in aging brains. PRM-SRS imaging also revealed subcellular distributions of sphingosine and cardiolipin in the human brain sample. Compared with other techniques, PRM-SRS demonstrates unique advantages, including faster data processing and direct user-defined visualization with enhanced chemical specificity for distinguishing clinically relevant lipid subtypes in different organs and species. Our method has broad applications in multiplexed cell and tissue imaging.

## Introduction

Current lipidomic studies such as shotgun lipidomics can identify different lipid subtypes with high sensitivity and specificity. Albeit highly sensitive, such methods rely on mass spectrometry (MS), nuclear magnetic resonance (NMR), or other techniques that are destructive to cells and tissues^1–4^. Conventional matrix-assisted laser desorption/ionization MALDI-MS imaging enables label-free lipid imaging but it has a lateral resolution on the order of cell diameters (∼10 µm) and destroys the sample during the process. In addition, 3D MALDI-MS imaging relies on serial sectioning of the sample, and the lipid species that are resolvable are limited to those with the highest ion yields. Other optical techniques have been developed to non-destructively visualize spatial distributions of lipid pools^5^ as well as metabolic flux^6^ at subcellular resolution, but they rely on markers, such as fluorescently labeled antibodies and transfected biosensors, which may alter the native distribution of lipids in cells or tissues. It is difficult to use labeled optical imaging to differentiate diverse molecular species simultaneously, since the diversity of lipid species far exceeds the specificity and availability of optical tags and dyes. Therefore, label-free optical imaging is instrumental. Stimulated Raman scattering (SRS) microscopy has demonstrated advantages of non-destructive 3D imaging with subcellular resolution in a label-free manner^7,8^. Recent work has even demonstrated quantitative mass concentration measurements of lipids, proteins, and water^9^. For label-free SRS imaging microscopy, the chemical specificity is achieved through hyperspectral imaging (HSI) or training of a deep learning model^10^. Lock-in free multiplex SRS imaging can rapidly extract hundreds of morphological or metabolic features in situ to understand lipid metabolism in cancer cells^11^. Despite these advancements, there has been no report on distinguishing multiple lipid subtypes in cells and tissue samples by using nondestructive label-free optical imaging methods.

In addition to imaging technologies, post-processing methods/algorithms also contribute to producing high resolution images. Recent work on Raman HSI analysis using multivariate curve resolution alternating least squares (MCR-ALS) algorithm has demonstrated effective deconvolution of chemical species without disturbing the native distribution of biomolecules^12^. However, a higher spectral resolution may entail prohibitively long imaging time. In such HIS, unmixing lipid species using unsupervised methods can be computationally expensive and lack the ability to directly identify a chemical species without manual association *posteriori*. For example, the MCR-ALS approach converts a complex spectrum to a linear combination of component spectra, but it can take 30 minutes to process a 512 × 512-pixel hyperspectral image and presupposes the number of chemical species in a sample. The result displays a pixel’s identity by their relative proportional composition of reference species. However, this is rarely feasible in a complex biological sample. Singular Value Decomposition (SVD) can estimate the number of components, but analytical results may be sensitive to slight deviations from the exact number of components. Clustering and segmentation of image pixels may be informed by MCR-ALS, but the precise molecular identities of the highlighted pixels may still be unknown, as there is no guarantee that the unmixed components correspond to a lipid subtype.

Spectral reference matching approaches, also known as spectral angle mapping, have been widely applied to Raman spectra analyses by quantifying the spectral similarity between an image pixel spectrum and a known reference spectrum^13^. **Figure 1** shows the general process of reference matching approach applied to hyperspectral imaging. First, spectra of the target analytes in the image are acquired using spontaneous Raman spectroscopy (**Fig. 1A**) and preprocessed for background removal and normalization (**Fig. 1B**). Hyperspectral Raman microscopy imaging is next performed on the sample of interest, in which each pixel contains specific spectral information (**Fig. 1C-D**). Then each pixel’s spectrum is preprocessed in the same way as the reference spectrum (**Fig. 1E**) and is analyzed with respect to the reference spectrum by calculating the cosine similarity score (**Fig. 1F**).

**Fig. 1.**
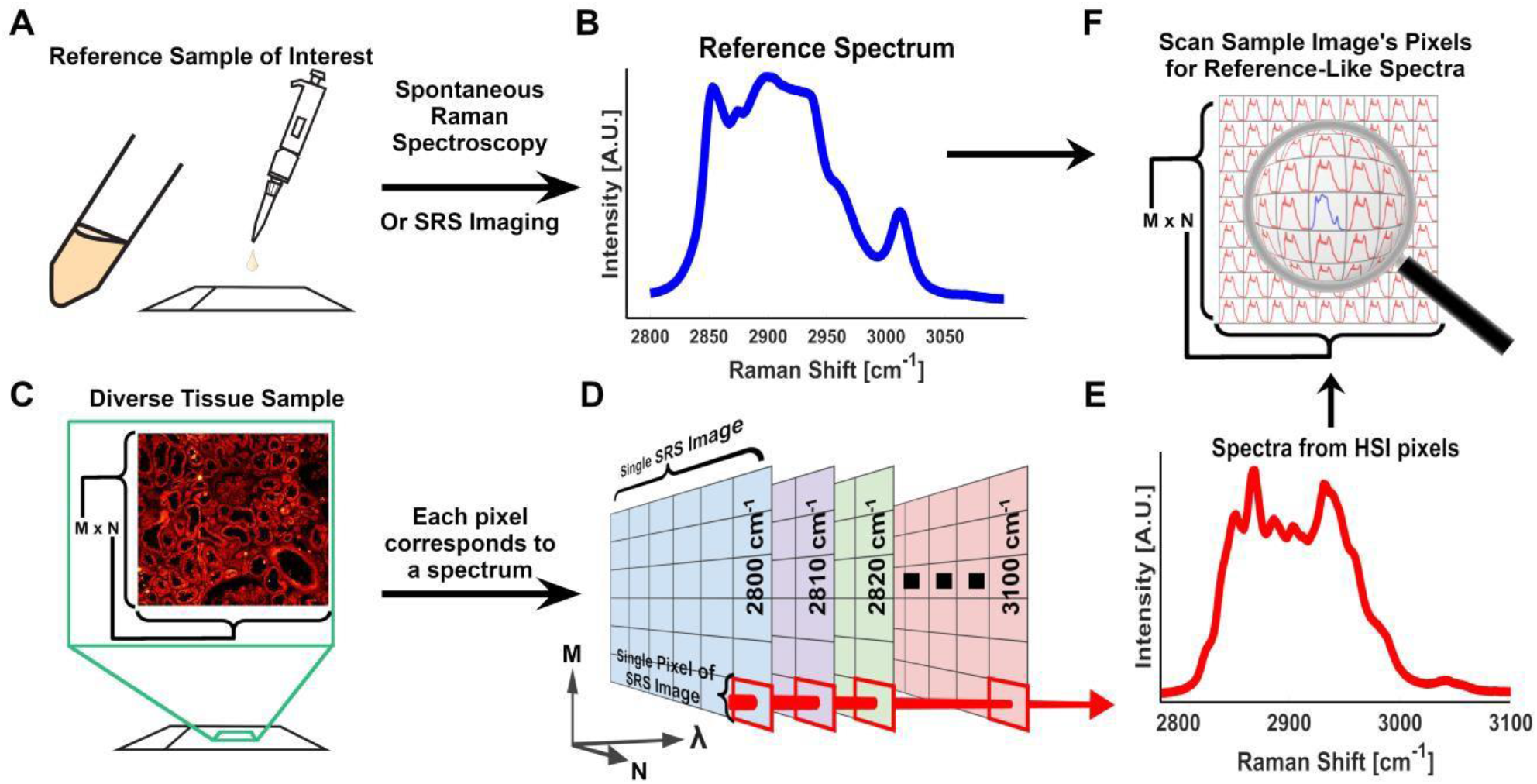
General Reference Matching Method. **(A)** A reference spectrum of a lipid subtype standard is acquired by spontaneous Raman spectroscopy and **(B)** preprocessed to normalize and remove background. **(C)** A sample is imaged using SRS to generate **(D)** a HSI. **(E)** Each pixel of the HSI is a vector of intensity values that represent the Raman spectrum at that pixel. **(F)** Similarity scores are calculated between each pixel and the reference spectra.

However, spectral reference matching approach has low specificity, and the high incidence of false positives has made it difficult to implement in vibrational spectroscopy. This is due to the peak position and intensity differences among various equipment produce uncertainty that overshadows the subtle differences between lipid subtypes. To enhance the specificity for accurately distinguishing different lipid species, we developed a Penalized Reference Matching (PRM) algorithm and applied it to SRS (PRM-SRS) microscopy to improve imaging contrast, with a breadth to accommodate a diverse library of more than 30 lipid subtypes. This method is extremely fast and can process a 512 × 512 pixel hyperspectral image stack within one minute. In this study, we demonstrate broad applications of PRM-SRS in differentiating lipid subtypes and mapping their spatial distributions in cells and tissues. This new method will provide quantitative and qualitative insights into different roles of lipid species in multiple biological processes.

## Results

### Developing a Penalized Reference Matching Method

SRS HSI pixel intensities were interpolated because reference matching by Euclidean distances implies that both the reference and observed spectra share the same number of data points. After all spectra were adjusted to the same interpolated resolution, they were simplex normalized using equation 1,

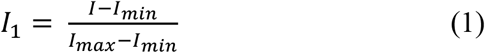

where *I*_*min*_ is the minimum value and *I*_*max*_ is the maximum value in the pixel spectrum I. The normalized pixel spectra *I*_1_ were then divided by their Euclidean norm as shown in equation 2

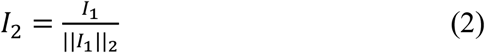

where *I*_*2*_ is the normalized signal of the pixel spectrum. Reference spectra from spontaneous Raman acquisitions follow the same pre-processing steps as the HSI pixel spectra, as shown in equations 3 and 4 below.

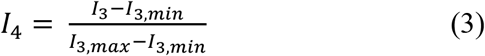

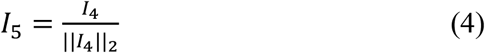

where *l*_3_denotes spontaneous Raman spectra, and *I*_*5*_ is the interpolated signal of the reference spectrum. Due to the nature of Raman spectral intensity, similarity scores between each pixel spectrum and the reference spectrum were calculated using the dot product of *I*_*2*_ and *I*_*5*_.

These spectroscopic methods have been deployed for several decades. However, due to the high incidence of false positives, direct label-free characterization of multiple lipid subtypes in cells and tissues has not been achieved in optical imaging. To address this issue, we added a penalty term to the canonical cosine similarity algorithm, which decreases the false positive rates by proportionally reducing the similarity score with the positional discrepancy to the best spectral match (**Fig. 2A-C**). This process is summarized as

**Fig. 2.**
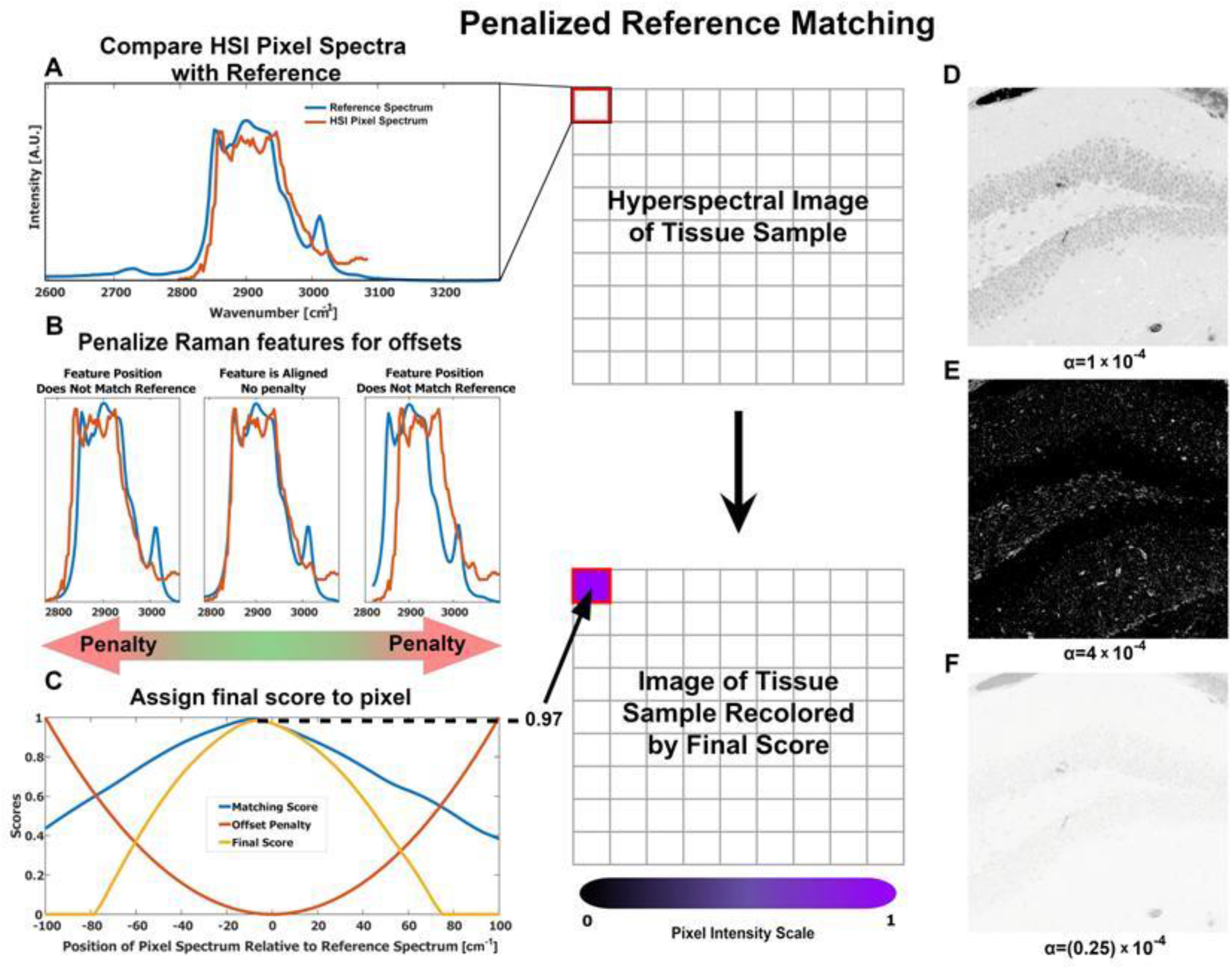
Penalized Reference Matching Method. **(A)** A lipid reference signal (blue) was collected by spontaneous Raman spectroscopy, and SRS HIS pixel spectrum (orange) was compared with the reference spectrum. (**B)** Example demonstration of shifts in pixel’s spectrum: 7.13 cm^-1^ left shift and 21.39 cm^-1^ right shift. These positional offsets decrease the final similarity score because the penalty term scales exponentially to the positional offset. (**C)** Matching scores were retrieved by dot product between the reference signal and SRS pixel’s signal. Then the penalty - estimated with a quadratic function – is subtracted from each matching score. When the shift wavenumber is high, a high penalty will be given. The highest value in this score curve will be used as the similarity score between this pixel spectra and that of the pure reference standard. **(D)** Image illustrating the distribution of cardiolipin in a murine dentate gyrus sample with a penalty coefficient of α = 1x 10^−4^. **(E)** Image with the penalty coefficient α = 4×10^−4^. The full range of the lipid reference was not clearly shown due to over-penalization. **(F)** An over-saturated image with penalty coefficient α = 0.25×10^−4^. Almost all pixels have a high similarity score due to under-penalization.

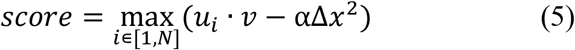

where ***u*** represents the interpolated signal of a pixel’s shifted spectrum at various positions; *v* represents the interpolated signal of the reference spectrum; *α* is the penalty coefficient; Δ*x* is the deviation in position of the spectrum in ***u*** from the initial observed position; and *N* is the number of interpolated signals, which depends on the spectral resolution of the HSI. The penalty term *α*Δ*x*^2^ inherently addresses the slight positional deviations due to the diverse chemical environment, as well as the variations in instrumentation such as thermoelectric noise, lensing, and other interference. Without this term, even if the spectral shape of a pixel matches the reference spectrum (**Fig. 2A**), the final similarity score may still be low when the positions of the peaks differ greatly (**Figs. 2B-C**). With the penalty term, all pixel spectra are evaluated as if they occur at multiple Raman shifts, and the highest similarity score is returned.

By leveraging positional information in addition to peak amplitude shape, the breadth of similarity score is increased, akin to an increase in contrast (**Fig. 2D-F**). This ensures that pixels with similar shapes and positions are scored consistently, enhancing the specificity of assignment of chemical identity.

Most images collected in this study were taken from the Raman CH stretching region (2700 cm^-1^ to 3150 cm^-1^) with 75 total Raman shifts (a spectral resolution of 5.6 cm^-1^). The position deviation Δ*x* was the shift of peaks in the spectrum. We assessed several values for the penalty coefficient and chose α =1×10^−4^. At a higher value (4×10^−4^), the image contrast was to too high to show the full dynamic range of signals, whereas a lower α (0.25×10^−4^) caused over-saturation in images (**Fig. 2D-F**).

### Mapping cholesterol levels in Drosophila fat body using reference spectra

As a proof of concept, we first applied the PRM algorithm to detecting and comparing cholesterol levels in fat body tissues of young and old Drosophila. Analogous to mammalian liver and adipose tissue, Drosophila fat body has been used extensively to study lipid metabolism. We collected fat body tissue spectra from young and old flies using spontaneous Raman spectroscopy and compared them to reference spectra of cholesterol at the fingerprint (750 cm^-1^ to 1650 cm^-1^) and CH-stretching (2700 cm^-1^ to 3150 cm^-1^) regions (**Fig. 3A-D**). Compared to samples from young flies, fat body samples from old flies showed significantly higher similarity scores to the cholesterol reference spectra in both regions (**Fig. 3E-F**), indicating elevated cholesterol content in old flies. This result is consistent with the published data ^14^. This analysis demonstrates our PRM algorithm as an effective method for rapid lipid mapping in tissues in situ.

**Fig. 3.**
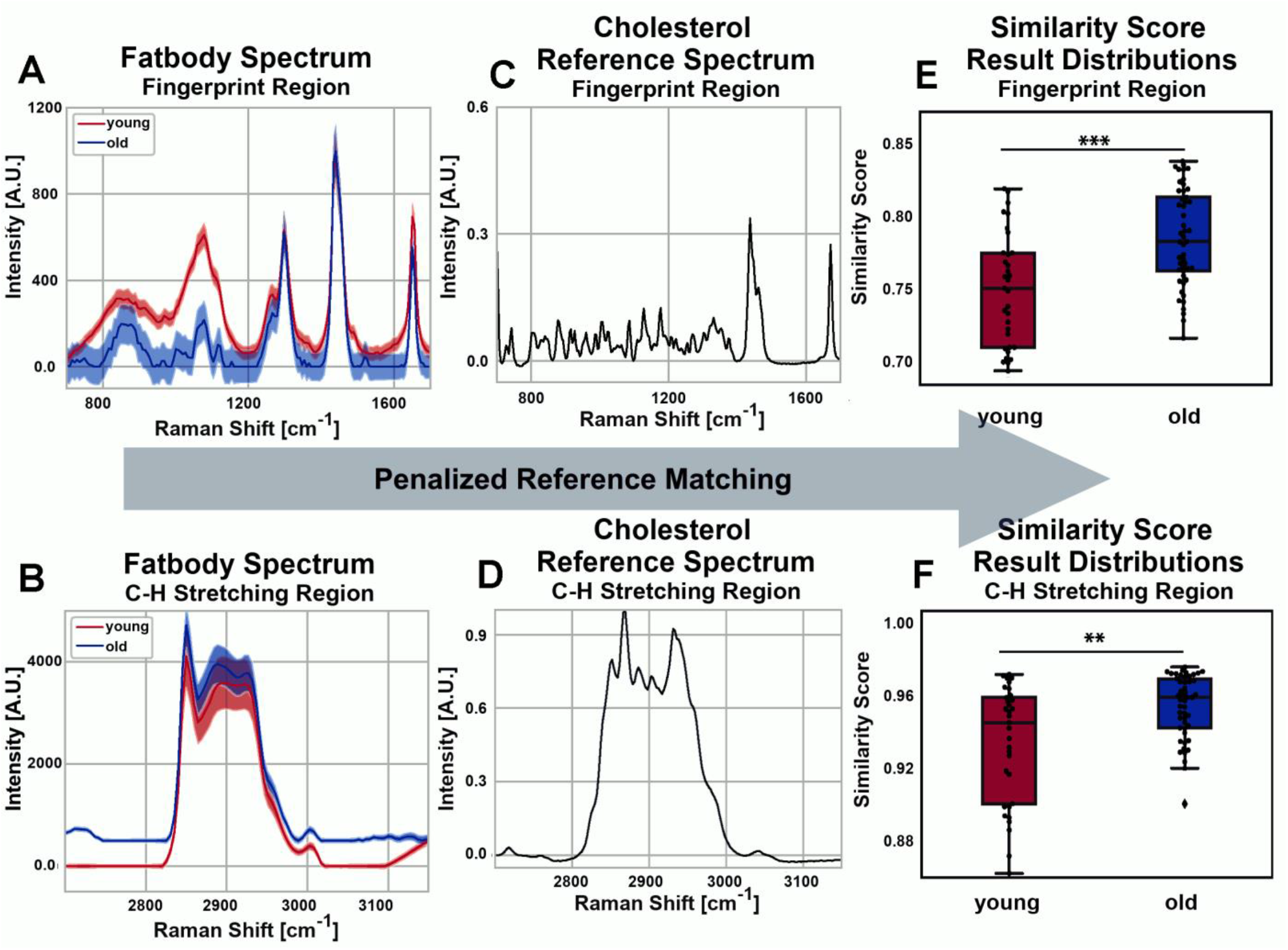
Spectral PRM cross correlation in the fingerprint and CH stretching regions. **(A, B)** Raman signals from fat body tissues of young and old Drosophila in the fingerprint and CH regions. (**C, D)** Raman signals of the cholesterol reference standard in fingerprint and CH stretching regions, respectively. (**E)** Fingerprint region similarity scores of young and old Drosophila fat body samples to the cholesterol reference standard. p = 4.85×10^−4^ by Wilcoxon rank sum test. (**F)** CH-stretch similarity scores of young and old Drosophila fat body samples to the cholesterol reference standard. p = 0.0037 by Wilcoxon rank sum test.**p<0.01, ***p<0.001.

Depending on the biological questions to address and Raman scattering equipment available, either the CH-stretching or fingerprint region in a Raman spectrum may be the focus of a study. Both the regions can be used to analyze changes in biomolecule distribution, pathological structures (such as amyloid plaques) and other morphological characteristics^15–19^. Although both spectral regions achieved similar results, the fingerprint region generated more statistically more significant results, with a rejection level of p<0.001. This is likely because the fingerprint region contained more definitive features, and the CH-stretching region spectra possessed low intensity shoulders below 2800 cm^-1^ and above 3000 cm^-1^ that may lead to a higher similarity score between samples since both spectral data sets matched in those regions where the intensity was zero. Importantly, this demonstrates that similarity scores generated from spectra data by PRM-SRS can be used to estimate the level of biomolecules in the samples.

### Using PRM-SRS to detect cardiolipin changes in cells

After validating the efficacy and robustness of PRM in spontaneous Raman spectral analyses, we next extended the algorithm to analyzing stimulated Raman scattering (SRS) images. To evaluate the spatial accuracy and quantitative approximation of PRM-SRS, we first benchmarked it against fluorescence microscopy images. Using PRM-SRS imaging, we examined cardiolipin (CL), an essential phospholipid in the inner mitochondrial membrane, in cultured HEK293 cells. CL is synthesized in the inner mitochondrial membrane in consecutive reactions catalyzed by enzymes, including phosphatidylglycerophosphate synthase 1 (PGS1), phosphatidylglycerophosphate phosphatase (PTPMT1) and cardiolipin synthase (CLS1)^20,21^. PGS1 is essential for CL synthesis, and expression of an enzyme-deficient mutant PGS1 leads to a reduction of PGP (Phosphatidylglycerophosphate) and CL in CHO cells^22^. We generated stable HEK293 cell lines with downregulated PGS1 (shPGS1). PGS1 down-regulation was confirmed by immunofluorescence analysis using a PGS1-specific antibody (**Supplementary Fig. S1**). Following staining with nonyl acridine orange (NAO), a fluorescent dye with high affinity for CL^23^, cells were analyzed using both two-photon fluorescence (TPF) microscopy and PRM-SRS. To demonstrate the specificity of SRS signals for CL, we compared control cells with shPGS1 cells. PRM-SRS analysis of the hyperspectral images was consistent with TPF images in both control and shPGS1 cells (**Figs. 4 A-B**). Quantitative analyses of SRS images and fluorescence images both showed significant decreases of CL signals in shPGS1 cells compared with control cells (**Figs. 4 C-D**). These results demonstrate the ability of PRM-SRS to quantitatively detect CL changes in cells, and its potential for visualizing lipid metabolic dynamics at the subcellular scale.

**Fig. 4.**
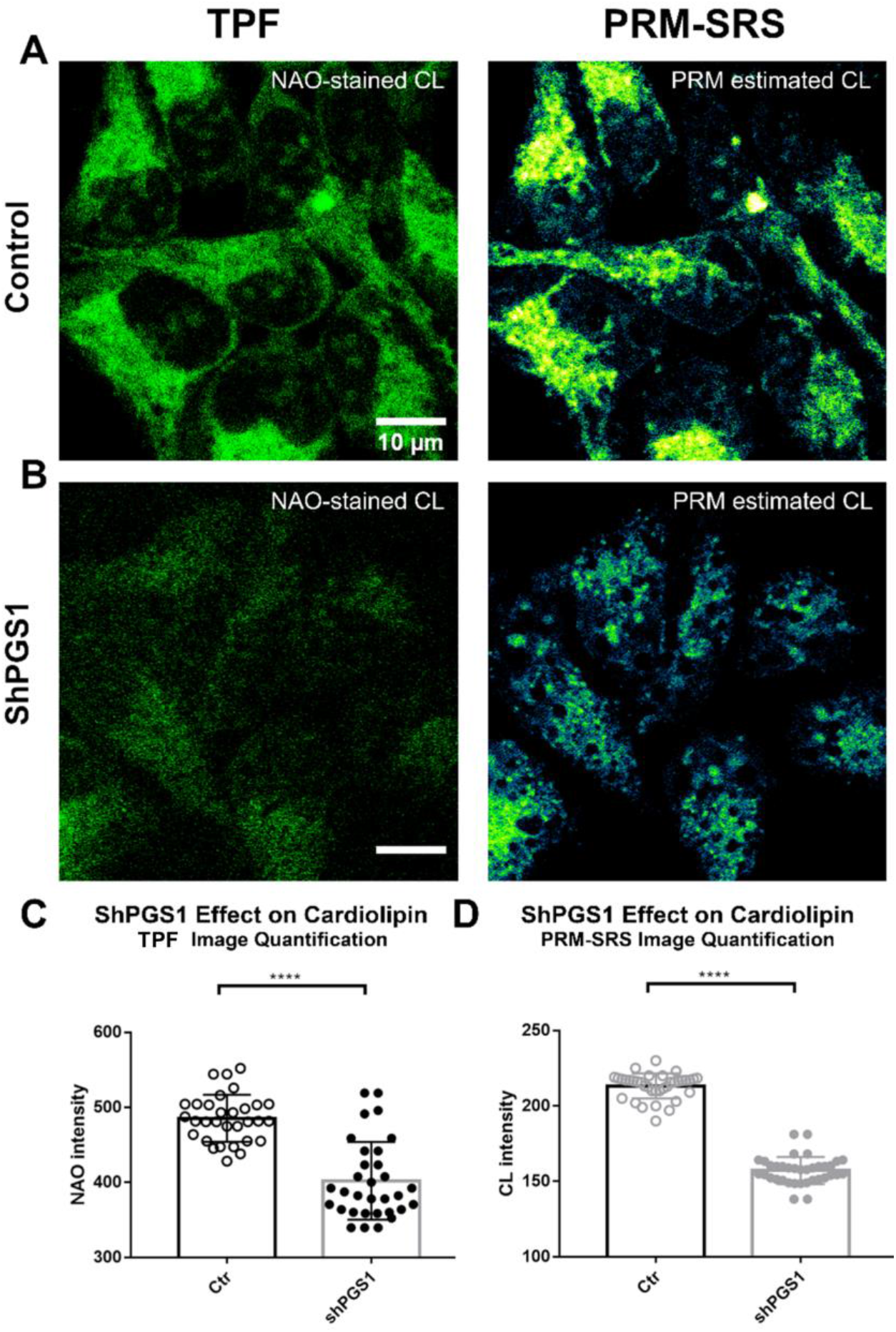
PRM-SRS and fluorescence staining show similar results.**A, B)** Comparison **of PRM-SRS and fluorescence staining in control cells expressing** shCtr (Ctr; **A**) and PGS1 knockdown (shPGS1; **B**) HEK293 cells. Panels on the left, two photon fluorescence microscopy (TPF) images following nonyl acridine orange (NAO)-labeling of CL. Panels on the right, label-free SRS hyperspectral images of CL at the CH-stretching region. Quantitative analyses of NAO staining signal intensity (**C**) and PRM-SRS imaging signal intensity (**D)** of CL in control and shPGS1 cells. Significantly decreased signals in shPGS1 cells were detected by both TPF and PRM-SRS microscopy. Data are presented as mean ± SEM and analyzed by One-way ANOVA with Bonferroni post hoc test; ****p<0.0001. Scale bar, 10 µm.

### PRM-SRS tracking clinically relevant lipid subtype biomarkers in human kidney tissue

We then applied PRM-SRS to characterizing lipid subtypes in human kidney tissue, a structurally and functionally complex tissue composed of more than 50 cell types^24^. Cholesterol, ceramides, and triacylglycerides are among the most abundant lipid species in the kidney. Dyslipidemia is frequently observed in nephrotic syndrome (NS) and each stage of chronic kidney disease (CKD)^25^. The glomerulus, the filtration unit of the nephron, is a network of capillaries that sequesters lipid species as an initial step of filtration and is decorated with lipid droplets. Wrapping around the capillary of the glomerular tuft are podocytes, making up the visceral epithelial lining of the Bowman’s capsule. We used healthy reference human kidney tissue sections to showcase the application of our PRM-SRS in imaging different lipid subtypes in structurally complex tissue samples.

In formalin-fixed kidney tissue sections, SRS imaging detected overall lipid distribution in the morphologically distinct structures, such as glomeruli, tubules and blood vessels (**Figs. 5A, 5B**). Using PRM-SRS, we observed lipid distribution and estimated relative concentrations of lipids in different structure, such as lipid droplets in podocytes and eosinophilic bodies near tubules (**Figs. 5 A-B)**. PRM-SRS imaging revealed distribution of distinct lipid subtypes in the glomerulus and surrounding structures in situ, including triacylglycerol (TAG), cholesterol, cholesterol ester and C12 ceramide, with the 90^th^ percentile similarity scores to the corresponding pure lipid reference spectra (**Fig. 5C**).

**Fig. 5.**
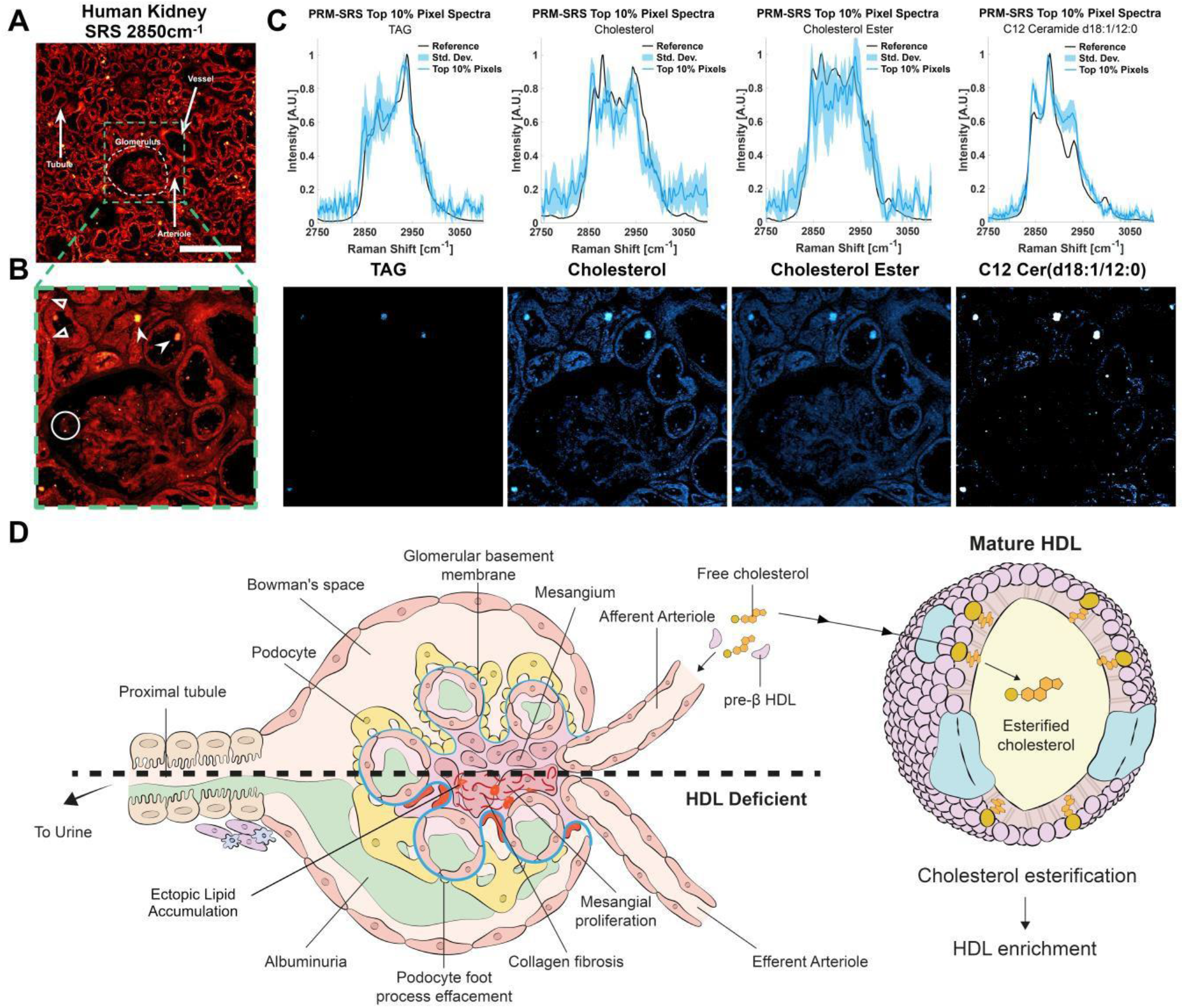
Label-free hyperspectral detection of different lipid subtypes *in situ* using PRM-SRS. **(A-B)** SRS image of a human kidney tissue section at 2850 cm^-1^. Panel B shows the enlarged image of the boxed area in panel A. Hollow arrowheads, intracellular lipid droplets in tubules. Solid arrowheads, eosinophilic bodies. Circles, lipid droplets sequestered by podocytes in the glomerulus. Scale bar, 200μm. **(C)** PRM-SRS spectra (top panels) and images (bottom panels) of different lipid subtypes of interest show the distribution of the similarity scores, each with the same contrast levels. Spectra of the top 10% of similarity score pixels overlaid on the reference spectrum for each lipid subtype show consistent matches. **(D)** Schematic diagram of glomerular pathologies associated with dyslipidemia in kidney diseases. (adapted from reference^49^ Reidy et al, 2014 10.1172/JCI72271 with permission) Scale bar, 200 µm.

Dyslipidemia is manifested as elevated levels of serum triglycerides, cholesterol, and very low to intermediate density lipoproteins. Common initial abnormalities include decreased production and activity of lecithin-cholesterol acyltransferase which decreases high-density lipoprotein (HDL) levels and maturation of HDL cholesterol^26^. The regulation of HDL cholesterol is tightly controlled by several organs, but generally entails the esterification of cholesterol into cholesterol esters, which move towards the center of HDL particles, along with neutral TAG lipids. This maintains a favorable cholesterol gradient as these HDL particles become enriched by sequestering cholesterol and fatty acids from other lipoproteins. Although mature lipoproteins are too large to pass the glomerular filtration barrier, lipids and lipid-bound proteins from lipoproteins may affect overall renal lipid metabolism^27^. Our ratiometric imaging revealed that there is a greater amount of non-esterified cholesterol in the lipid particles than neighboring structures. These pools of cholesterol may represent those yet to be enriched or ectopic deposits (**Supplementary Fig. S2**). Ceramides are also abundant in the kidney and play a crucial role in regulating cellular processes by binding cholesterol and other lipoproteins^28^. Ceramides, e.g., C12 ceramide, show high similarity with pixel spectra in lipid droplets and lipoprotein particles (**Fig. 5C**). In nephropathies, ectopic lipid deposits in the glomerular mesangium and proximal tubules are typically concurrent with low HDL levels^26^. Other characteristics of glomerular nephropathies are depicted in **Fig. 5D**. The ability of PRM-SRS to track the lipidomic profile in tissues collected from patients at various stages of diseases will generate critical data for changes in these macromolecules over time, and with associated biological variables. Such studies will provide insights into assessing severity, progression or prognosis of various lipid metabolic diseases.

### Mapping lipid subtype distributions in Drosophila fat body

In addition to the detection of lipid subtypes, PRM-SRS can also provide information on subcellular distribution, including co-localization, of different lipid subtypes. We visualized lipids in Drosophila fat body cells (**Fig. 6**) using the reference spectra of phosphatidylcholine (PC) and phosphatidylethanolamine (PE). PC and PE are the most prominent lipid subtypes in cell/organelle membranes and can be quantitatively distinguished from each other. Even without ratiometric analysis or additional experimental groups, PRM-SRS images clearly demonstrate the distributions of PE and PC on the lipid droplet membrane relative to the total lipid content (**Fig. 6**). To our knowledge, these data have demonstrated, for the first time, that multiple lipid subtypes can be measured from a single HSI stack in multiplex SRS imaging.

**Fig. 6.**
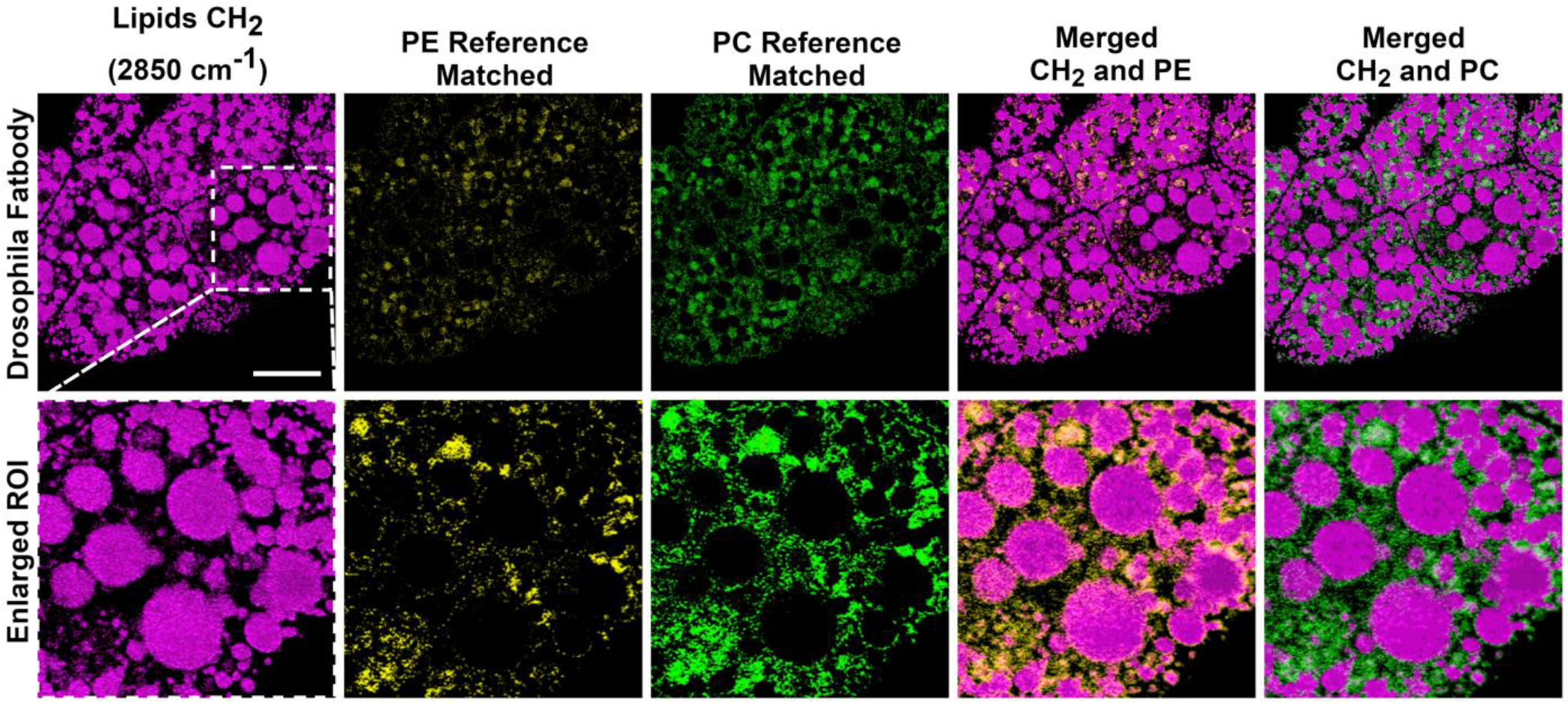
Membrane lipid subtypes co-localized with the lipid droplet surface in Drosophila fat body cells. PRM-SRS images of total lipids (CH_2_ stretching), PE, and PC show that the membrane lipid species are readily visible near the lipid droplet surface and in the interstitial space. Scale bar, 20 µm.

### Analyzing lipid subtypes in mouse brain samples

We next applied PRM-SRS to analyze lipid metabolism in the context of the aging using mouse hippocampal samples. We visualized and compared cholesterol, PC, and PE levels in hippocampal samples from young (3 months) and old (18 months) mice (**Figs. 7 A-C; F-G**). We also generated ratiometric images for quantitative analysis, since the signal intensity has a linear relationship with the concentration of chemical bonds of the molecules detected. Ratiometric imaging analyses showed increased Cholesterol/PE ratio in subregions of granule cell nuclei (**Fig. 7D, 7I**; red circles). This increase in the Cholesterol/PE ratio was more prominent and detected in more granule cells in the old brain samples as compared with the young ones (compare **Fig. 7D** with **7I**), demonstrating altered cholesterol and/or PE metabolism in the old brains. These results show that ratiometric PRM-SRS imaging can detect changes in differential spatial distribution of various lipid subtypes even when such changes are not obvious in images of individual lipid subtypes.

**Fig. 7.**
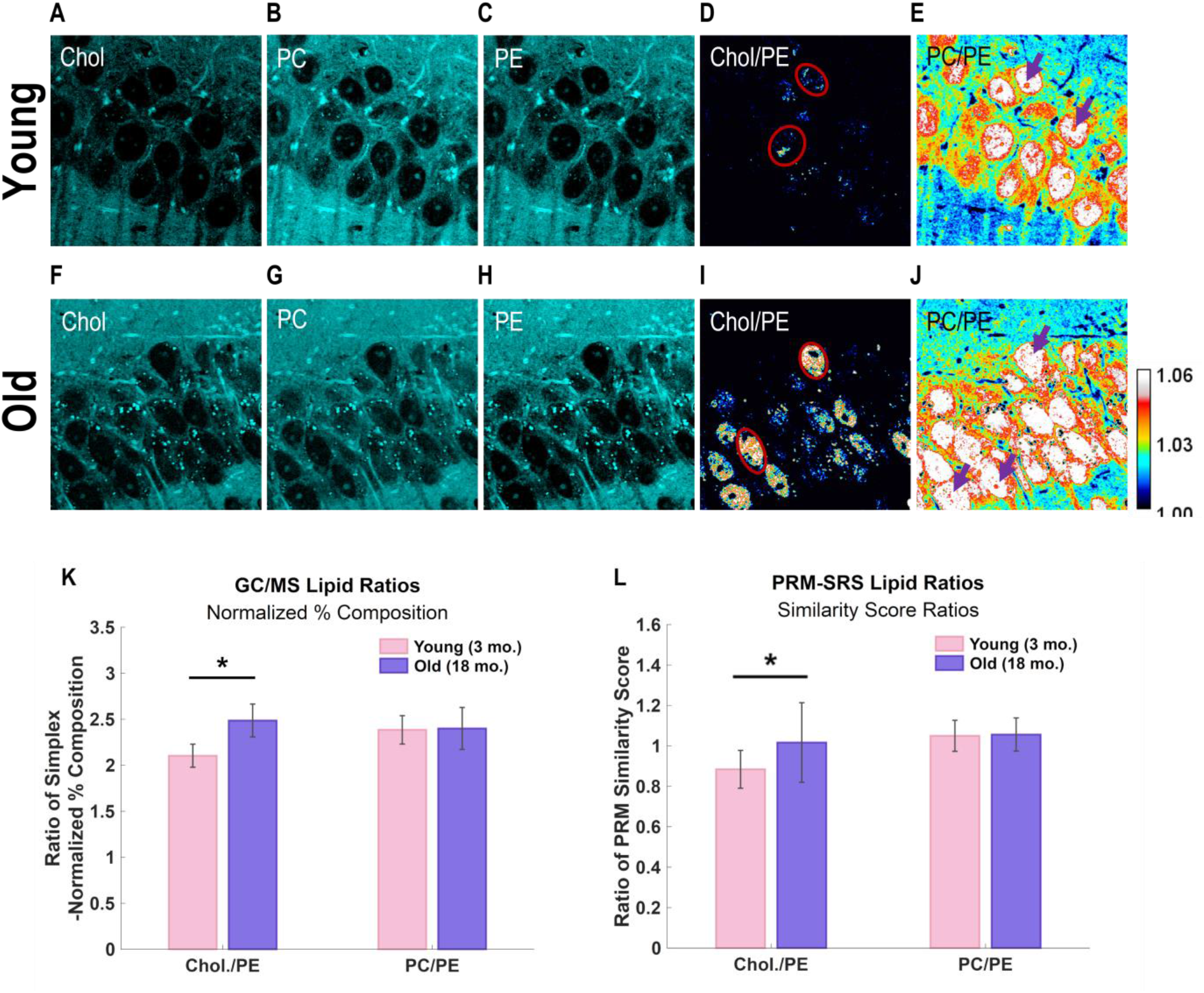
PRM-SRS imaging of mouse hippocampal samples. **(A-J)** PRM-SRS hyperspectral detection of cholesterol, PC, and PE in hippocampus samples from young and old mice. Overall intensity of detected lipid subtypes shows distinct patterns, with old brains showing higher cholesterol to PE ratio, but relatively consistent levels of PE and PC. **(D, I)** Ratiometric images of cholesterol to PE shows more nuclei with higher cholesterol/PE ratio in the old brains. Selected nuclei are marked by red circles **(E, J)** Ratiometric images of PC relative to PE show higher PC/PE ratio in granule cell nuclei of both young and old brains, but the spatial distribution of the ratio is more heterogeneous in young samples (see nuclei marked by purple arrows). **(K, L)** Mass spectrometry (K) shows results consistent with that obtained by ratiometric PRM-SRS imaging (**L**). Summed concentrations of lipids were simplex normalized and displayed in the ratio form. Error bars represent standard deviation. **(L)** Ratiometric image intensities, corresponding to the ratio of PRM similarity scores of lipid subtypes, are plotted. Error bars represent standard deviation. Scale bar, 20µm.

Ratiometric images of PC/PE showed higher levels of PC relative to PE in the granule cell nuclei of the dentate gyrus in both young and old mice, but lower levels outside the nuclear regions (**Fig. 7 E, J**). Compared to the young brain sample, the old brain sample showed no significant changes in the average PC or PE levels in the granule cells both in the individual imaging channels (**Fig. 7 B,C,G,H**), and the ratiometric images (**Fig. 7 E,J**). This is consistent with the results from Gas Chromatography Mass Spectrometry (GC-MS) (see **Fig. 7K, L**) results from mice hippocampus. However, we noticed spatial distribution differences in the PC to PE ratio between young and old samples. The ratiometric images (**Fig. 7 E,J**), reveal that more granule cell nuclei had uniformly higher PC/PE ratio in the old brain sample, whereas the nuclei in the young sample showed less even distribution of the PC/PE ratio (red areas; see those nuclei marked by purple arrows) (**Fig. 7 E,J**). These data suggest altered synthesis, accumulation or clearance of PC and/or PE in the granule cells in the old brains, consistent with a previous report^29^. Since PE is a precursor of PC, higher PC to PE ratios inside the older hippocampal granule cells suggest that aging brains may have altered CTP:phosphocholinecytidylyl transferase (CCT) activity – a rate limiting PC synthesis enzyme with a predominantly nuclear localization ^30^. This finding is significant because both PRM-SRS imaging and GC-MS analysis show that the younger brain samples contain less cholesterol relative to PE than old ones. However, only through ratiometric analysis we were able to detect differential subcellular distribution of lipids, including cholesterol/PE and PC/PE ratio in the nuclei (**Fig. 7D, 7I; 7E, 7J**).

For comparison, we analyzed the same samples using GC-MS to quantify cholesterol/PE and PC/PE ratios (**Fig. 7K**). The PRM-SRS images of nuclei in the tissues were manually segmented using ImageJ for quantification of cholesterol/PE and PC/PE ratios (**Fig. 7L**). Ratiometric PRM-SRS imaging analysis results are consistent with that obtained by traditional GC-MS. These data suggest that PRM-SRS can be used for quantitative lipidomic imaging analyses in tissue samples in the future.

### Detecting lipid subtype distributions in human brain tissues

Sphingosine is another crucial lipid subtype whose metabolic alteration has been suggested as a biomarker for neurodegenerative diseases, such as Alzheimer’s, Parkinson’s, and Huntington’s diseases^4,31^. To visualize individual cells, we used label-free optical SRS histology (SRH) imaging of human brain sample (See supplementary Fig. S3) to create virtual histology images similar to that of hematoxylin-and-eosin (H&E) staining as previously reported^35^. Using PRM-SRS, we visualized sphingosine and CL simultaneously in the human brain tissue sections (**Fig. 8 A, 8B**). Superimposition of sphingosine and CL images illustrates their relative distribution in brain cells (**Fig. 8C, 8D**). Ratiometric imaging (**Fig. 8E**) and quantitative analyses (**Fig. 8F**) demonstrated a clear reduction in the CL to sphingosine ratio inside the nucleus, consistent with the fact that CL is mainly localized at the inner mitochondrial membrane but not in the nucleus. These results show that PRM-SRS can be used to visualize subcellular distribution of different lipid subtypes.

**Fig. 8.**
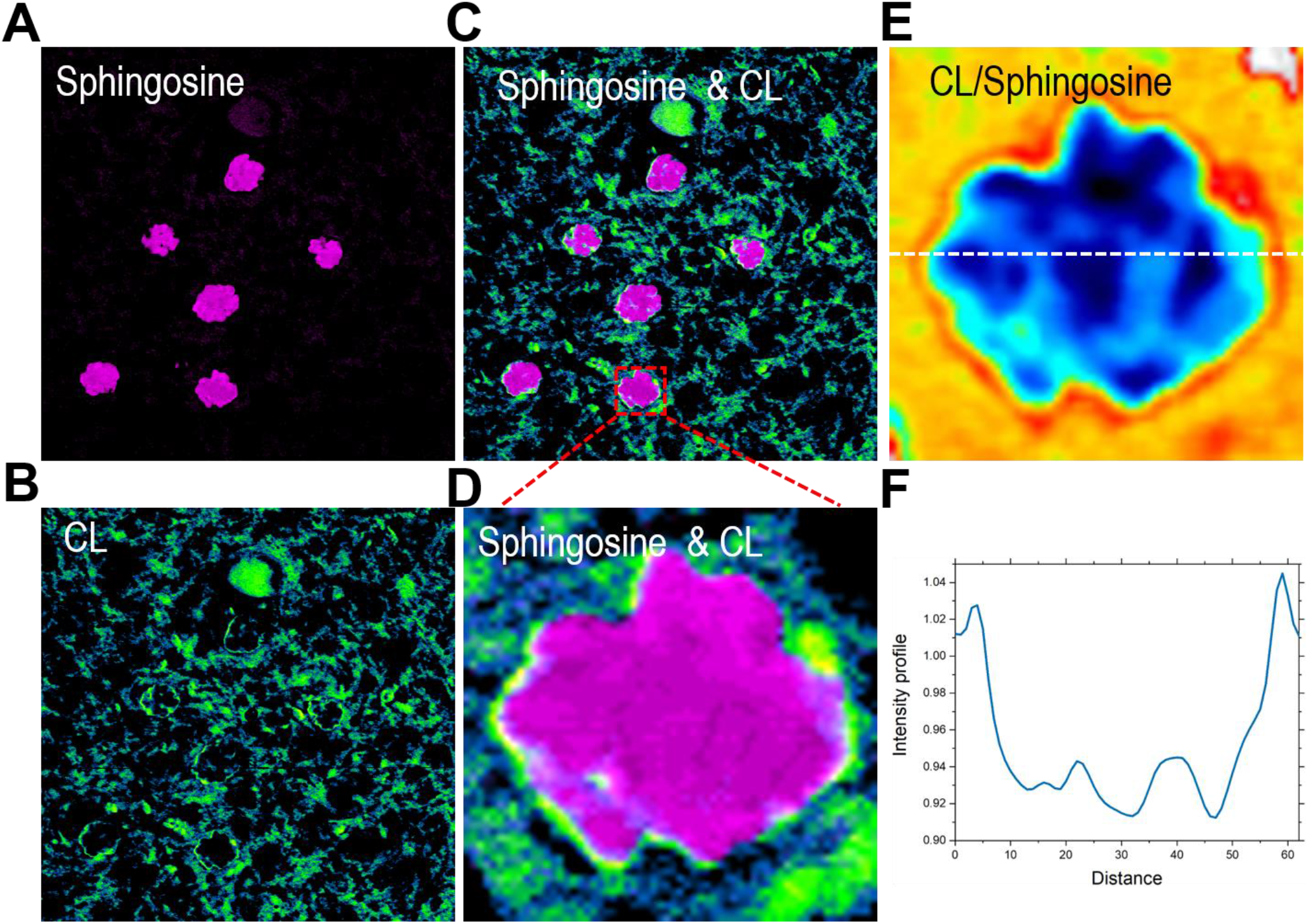
Hyperspectral SRS imaging detection of cardiolipin and sphingosine signals in a human brain tissue section. **(A)** Sphingosine signals in the brain tissue; **(B)** CL in the same region of interest; **(C)** merged image of CL and Sphingosine; **(D)** Zoomed-in image of a single brain cell with CL and Sphingosine signals. **(E)** Ratiometric image of CL to Sphingosine signals. **(F)** Intensity profile of **(E)** along the indicated white dashed line. Scale bar,10 µm.

## Discussion

In this study, we developed a PRM algorithm that can efficiently unmix and distinguish a variety of lipid subtypes from single SRS HSI stacks. Compared with fluorescence imaging, our PRM-SRS platform shows advantages of multiplexed lipid subtype visualization from single label-free HSI sets. This also represents a significant expansion in applications compared with traditional SRS imaging, which often relies on detecting lipids in the CH-stretching mode at 2850 cm^-1^.

With an improved contrast mechanism, PRM-SRS imaging enables us to detect a large number of lipid subtypes. It can generate both co-localization and ratiometric data of individual lipid subtypes simultaneously by mapping their spatial distributions and quantifying their relative concentrations. We have established a library of over 30 lipid subtypes, all of which can be distinguished by PRM-SRS simultaneously. In this study, we selected 8 lipid subtypes for a proof of concept (**supplementary Fig. S4**). Analyses of human kidney tissue samples indicate that PRM-SRS can be used to identify different lipid subtypes associated with renal diseases, suggesting potential application of PRM-SRS in diagnosis and prognosis of these diseases, including those associated with dyslipidemia. These label-free biomarkers may be instrumental in early detection of kidney diseases by detecting and measuring relative levels of different lipid biomarkers without the need to stain biopsied samples or perform destructive imaging, especially on limited clinical samples. Analyses of Drosophila fat body samples show that PRM-SRS can be used effectively in mapping spatial distributions of membrane lipids at the subcellular scale. These results highlight the ability to selectively visualize multiple lipid subtypes in a single image with the ease and freedom akin to single-peak labeled imaging without the need to actually label them. Analyses of mouse brain tissues demonstrate the importance of measuring relative lipid concentrations through ratiometric imaging, which reveals regionally different concentrations of lipid subtypes that may not be readily apparent in single-channel images. Although lipid subtype levels are not revealed in absolute concentrations, their relative levels are consistent with results from other modalities such as mass spectrometry (MS). Analyses of mouse and human brain tissues illustrate the capability of PRM-SRS in quantitatively mapping and analyzing distribution of different lipid subtypes within single cells. These analyses also confirm the cross-applicability of the fingerprint and CH-stretching spectral regions for quantitative analyses. The brain is a lipid-rich organ. Lipid subtypes such as cholesterol and sphingolipids are important components of the brain. Alteration in lipid subcellular distribution and metabolism impact on brain cell function ad have been associated with neurological diseases. Our PRM-SRS imaging shows that sphingosine, a catabolic byproduct of sphingomyelin, has a predominantly nuclear localization. Nuclear sphingomyelinase and sphingosine kinases regulate the release of ceramides and sphingosine, as well as the conversion to sphingosine-1-phosphate. These processes regulate cell proliferation and cell death^36^. A previous study showed that sphingosine kinase may shift from a cytosolic to a nuclear localization in the brain samples from Alzheimer’s disease patients^37^. Development of new technologies in imaging distinct lipid subtypes and their metabolism will enhance our ability to investigate molecular mechanisms underlying different brain disorders.

Depending on the equipment used and the sample of interest, careful tuning of the penalty term in the PRM algorithm is necessary. At present, the PRM-SRS platform should be used in a well-controlled environment to limit external chemometric dimensions. This way, spectral signals are more likely from molecular subtypes in the samples, rather than from noises. Since Raman peak intensities are multiplexed in the sense that a specific peak shape may be influenced by multiple molecules, it is critical that molecular makeup is as consistent as possible when using PRM-SRS to determine relative concentrations of different molecular subtypes. PRM-SRS may also be used together with other chemometric methods, such as GC-MS, for cross validation, as the incidence of false positive may still be high. Finally, detection of the vast variety of lipid subtypes may require further improvement in unmixing methods and greater spectral resolution, as the lipid subtype reference library is expanded, and more lipid subtypes are further evaluated. With some adjustments, such as using different reference libraries, this PRM-SRS platform can be extended to analyzing other molecules, including proteins, nucleic acids, and even clinically relevant molecular complexes (such as protein aggregates or oligomers). Using heavy water (D2O) probed SRS (DO-SRS), metabolic imaging can also distinguish de novo synthesized new biomolecules, including lipids, protein, and DNA^38–40^, from old existing biomolecules at the subcellular resolution. This ever-expanding library of molecular subtype references may warrant broader spectral regions, including the fingerprint, CH-stretching, and O-H stretching regions to increase the chemometric dimensions. Integration of statistical denoising and regression methods will help increase the power of the matching score. Application of higher order signal manipulations such as digital derivatives and wavelet analyses, will enhance the ability to extract the most prominent as well as subtle but important features.

In summary, this study presents a new hyperspectral imaging platform - PRM-SRS - that allows for direct identification of multiple molecular species in situ with subcellular resolution and high chemical specificity by leveraging the cross-applicability of spectral reference libraries and HIS methods. This PRM-based method can be applied to various microscopy setups, such as SRS, FTIR, and spontaneous Raman scattering spectroscopy. Compared with existing HSI methods, PRM-SRS shows a much-enhanced speed and efficiency. With appropriate reference spectra established, PRM-SRS can be used to detect a wide range of different biomolecules. This new platform can also be applied to studying metabolism of diverse types of biomolecules in cell and tissue samples, which will be highly useful for investigating metabolic changes under different pathophysiological conditions. With its easy implementation, PRM-SRS can also be combined with high-throughput methods, such as microfluidic/nanofluidic devices and single-cell apparatuses, or with large-area HSI mapping methods. The application of deep learning algorithms such as DeepChem may improve the imaging speeds further by utilizing single-frame femtosecond SRS imaging^10^. Thus, PRM-SRS has great potentials in multiplex cell and tissue imaging with a broad spectrum of applications.

## Materials and methods

### Reference Lipid standards

Reference lipid standards were purchased from Sigma-Aldrich (St. Louis, MO) or Avanti Polar Lipids (Alabaster, Alabama): cardiolipin 14:0/14:0/14:0/14:0 (CL), <u>C12 ceramide,</u> ceramide 18:1;2/17:0 (Cer), diacylglycerol 17:0/17:0 (DAG), hexosylceramide18:1;2/12:0 (HexCer), lyso-phosphatidate 17:0 (LPA), lyso-phosphatidylcholine 12:0 (LPC), lyso-phosphatidylethanolamine 17:1 (LPE), lyso-phosphatidylglycerol 17:1 (LPG), lyso-phosphatidylinositol 17:1 (LPI), lyso-phosphatidylserine 17:1 (LPS), phosphatidate 17:0/17:0 (PA), phosphatidylcholine 17:0/17:0 (PC), phosphatidylethanolamine 17:0/17:0 (PE), phosphatidylglycerol 17:0/17:0 (PG), phosphatidylinositol 16:0/16:0 (PI), phosphatidylserine 17:0/17:0 (PS), cholesterol D6 (Chol), cholesterolester 20:0 (CE), sphingosine, sphingomyelin 18:1;2/12:0;0 (SM) and triacylglycerol 17:0/17:0/17:0 (TAG).

### Sample Preparation

#### HEK293 Cell Cultures

HEK293 based T-REx™293 cell line was obtained from Invitrogen). Cells were cultured in DMEM supplemented with 5% fetal bovine serum (FBS), 1% penicillin/streptomycin (Fisher Scientific, Waltham, MA).

The control shRNA construct (shCtr) was as previously described^41^. A shPGS1 construct was designed to express shRNA against PGS1 (target sequence: 5’-TCGGGTTCCATCCGTTTAAAT-3’) in the plasmid vector Tet-pLKO-puro (Vector Builder Inc) to specifically down-regulate expression of PGS1following induction with tetracycline (Tet). The control shCtr and shPGS1 constructs were transfected into T-REx™293 cells using lipofectamine (Invitrogen). Following transfection, cells were selected using puromycin (1ug/ml) and stably expressing cell clones were obtained. Following Tet treatment (1ug/ml; 48hrs), control shRNA and shPGS1 cells were prepared for immunostaining, NAO staining or SRS imaging. Cells were passaged at 80% confluence and plated on #1 thickness laminin coated coverglass (GG12-laminin, VWR). After allowing cells to adhere to the coverglass for 2 hours, cells were fixed using 4% v/v PFA for 15 min and stained with 100 nM NAO in the dark for 30 min. Cells were SRS imaged transmissively through #1 thickness coverglass.

Immunofluorescence staining was performed following our published protocol^41^ using a polyclonal rabbit anti-PGS1(Sigma-Aldrich, Cat# AV48896) and secondary antibody conjugated with Alexa-488 (Abcam, Cat# ab150081).

#### Human Kidney Tissue Preparation

De-identified human kidney tissue sections (30μm) were prepared from 4% v/v PFA-fixed frozen biopsy samples using a Compresstome (VF-210-0Z, Precisionary). Samples were imaged between 1mm thick glass slide and #1 thickness coverglass, submerged in 1x PBS. The kidney cortex was isolated for imaging.

#### Human Brain Tissue Preparation

De-identified post-mortem autopsy human brain sections (6 μm) were prepared from formalin-fixed and paraffin-embedded cortex tissue of control subject without detectable and neuropathology as previously published^42,43^. The sections were deparaffinized following a published protocol ^13^. Subsequent SRS imaging was conducted with the tissue sections sandwiched in PBS between 1mm thick glass slides and #1 thickness cover glass.

#### Mouse Brain Samples

The young (3 months) and aged (18 months) mice were euthanized with 5% isoflurane, and then perfused with 4% paraformaldehyde. The brains were harvested and fixed in 4% paraformaldehyde at 4°C for overnight. The fixed brains were washed with PBS and cut into 120-μm thickness slices with Vibratomes (Precisionary). The brain slices were placed in the center of a spacer and sandwiched between glass slides and coverslip for hyperspectral SRS imaging.

#### Drosophila Fat body Samples

Wild type (*w*^*1118*^ stock #5905) were originally obtained from the Bloomington Stock Center and have been maintained in the lab for several generations. Fat bodies were dissected from day 7 adult flies and fixed in 4% PFA (in 1xPBS) for 15 min. Samples were imaged immediately using SRS microscopy for hyperspectral imaging.

### Spontaneous Raman Spectroscopy

Spontaneous Raman scattering spectra were obtained by a confocal Raman microscope (XploRA PLUS, Horiba) equipped with a 532 nm diode laser source and 1800 lines/mm grating. The acquisition time is 30 seconds. The excitation power is ∼40 mW after passing through a 100x objective (MPLN100X, Olympus). Output spectra are background subtracted and vector and simplex normalized. The individual pure lipid reference was placed on glass slides for spontaneous Raman spectra measurement. All lipid subtype reference spectra were acquired in the same manner.

### Stimulated Raman Scattering Microscopy

An upright laser-scanning microscope (DIY multiphoton, Olympus) with a 25x water objective (XLPLN, WMP2, 1.05 NA, Olympus) was applied for near-IR throughput. Synchronized pulsed pump beam (tunable 720–990 nm wavelength, 5–6 ps pulse width, and 80 MHz repetition rate) and Stokes (wavelength at 1032nm, 6 ps pulse width, and 80MHz repetition rate) were supplied by a picoEmerald system (Applied Physics & Electronics) and coupled into the microscope. The pump and Stokes beams were collected in transmission by a high NA oil condenser (1.4 NA). A high O.D. shortpass filter (950nm, Thorlabs) was used that would completely block the Stokes beam and transmit the pump beam only onto a Si photodiode for detecting the stimulated Raman loss signal. The output current from the photodiode was terminated, filtered, and demodulated in X with a zero phase shift by a lock-in amplifier (HF2LI, Zurich Instruments) at 20MHz. The demodulated signal was fed into the FV3000 software module FV-OSR (Olympus) to form the image during laser scanning. All SRS images were obtained with a pixel dwell time 40 µs and a time constant of 30 µs. Laser power incident on the sample is approximately 40mW. Stimulated Raman histology was performed following a published protocol ^33^.

### Gas Chromatography Mass Spectrometry (GC-MS)

Hippocampal region slices (n=4 per group) from 3 month old and 18 month old mice were homogenized in ethanol/water1:1 (v/v) and the homogenate were sent to Lipotype GmbH (Dresden, Germany) for mass spectrometry-based lipid analysis^44^. Lipids were extracted using a two-step chloroform/methanol procedure^45^. Samples were spiked with internal lipid standard mixture containing: cardiolipin 14:0/14:0/14:0/14:0 (CL), ceramide 18:1;2/17:0 (Cer), diacylglycerol 17:0/17:0 (DAG), hexosylceramide18:1;2/12:0 (HexCer), lyso-phosphatidate 17:0 (LPA), lyso-phosphatidylcholine 12:0(LPC), lyso-phosphatidylethanolamine 17:1 (LPE), lyso-phosphatidylglycerol 17:1(LPG), lyso-phosphatidylinositol 17:1 (LPI), lyso-phosphatidylserine 17:1 (LPS), phosphatidate 17:0/17:0 (PA), phosphatidylcholine 17:0/17:0 (PC),phosphatidylethanolamine 17:0/17:0 (PE), phosphatidylglycerol 17:0/17:0 (PG),phosphatidylinositol 16:0/16:0 (PI), phosphatidylserine 17:0/17:0 (PS), cholesterolester 20:0 (CE), sphingomyelin 18:1;2/12:0;0 (SM), triacylglycerol 17:0/17:0/17:0(TAG) and cholesterol D6 (Chol). After extraction, the organic phase was transferred to an infusion plate and dried in a speed vacuum concentrator. First step dry extract was re-suspended in 7.5 mM ammonium acetate in chloroform/methanol/propanol (1:2:4, V:V:V) and 2nd step dry extract in 33% ethanol solution of methylamine in chloroform/methanol (0.003:5:1; V:V:V). All liquid handling steps were performed using Hamilton Robotics STARlet robotic platform with the Anti Droplet control feature for organic solvents pipetting.

Samples were analyzed by direct infusion on a QExactive mass spectrometer (ThermoScientific) equipped with a TriVersa NanoMate ion source (Advion Biosciences). Samples were analyzed in both positive and negative ion modes with a resolution of Rm/z=200=280000 for MS and Rm/z=200=17500 for MSMS experiments, in a single acquisition. MSMS was triggered by an inclusion list encompassing corresponding MS mass ranges scanned in 1 Da increments^46^. Both MS and MSMS data were combined to monitor CE, DAG and TAG ions as ammonium adducts; PC, PC O-, as acetate adducts; and CL, PA, PE, PE O-, PG, PI and PS as deprotonated anions. MS only was used to monitor LPA, LPE, LPE O-, LPI and LPS as deprotonated anions; Cer, HexCer, SM, LPC and LPC O-as acetate adducts and cholesterol as ammonium adduct of an acetylated derivative^47^. Data were analyzed with in-house developed lipid identification software based onLipidXplorer^48^. Data post-processing and normalization were performed using an in-house developed data management system. Only lipid identifications with a signal-to-noise ratio >5, and a signal intensity 5-fold higher than in corresponding blank control samples were considered for further data analysis.

### Data Analysis

#### Image Processing

SRS images were converted to unsigned 16 bit images via MATLAB, and were filtered using a morphological top-hat algorithm with 8 structuring elements, where appropriate. Unless used in ratiometric calculations, images for display were background subtracted using a sliding paraboloid with a radius of one tenth the image length. Intensity profiles and color maps were generated from ImageJ. All images within a figure use the same contrast unless specified otherwise. Ratiometric and overlaid images were created using the Image Calculator function and Overlay function, respectively, in ImageJ. Statistical analyses were performed using SPSS.

## Acknowledgements

We thank Drs. K. Zhang, C. Metallo, F. Liu, and G. Schmid-Schoenbein for helpful discussions. We acknowledge support from UCSD Startup funds, NIH U54 pilot grant 2U54CA132378, NIH 5R01NS111039, and Hellman Fellow Award. We are grateful for the support of the Washington University Kidney Translational Research Center (KTRC) for kidney samples and the HuBMAP grant U54HL145608. We thank Dr. E. Bigio and Dr. M-M. Mesulam from Mesulam Center for Cognitive Neurology and Alzheimer’s Disease (MCCNAD) for providing the de-identified autopsy brain samples; and MCCNAD is supported by NIH P30 AG013854.

## Author contributions

L.S. conceived the idea and designed the project; W.Z. developed PRM-SRS algorithm and coded it with help from L.S., Z.L., and A.F.; Y.L. and A.F. carried out the imaging experiments and collected data with the help from L.S. and Z.L.; A.F., L.S., H.J., Y.L., and Z.L. analyzed the image and generated the figures; J.Y.W. helped with project design; X.C. prepared part of the cell and tissue samples; S. J. contributed to data interpretations for human kidney samples. J. Y. and H.S. contributed to data interpretations for mouse brain samples. F.G. and D.S. conducted mass spectrometry measurement; A.F. and L.S. drafted and revised the manuscript with the input from all other authors.

