## supplement for "Multi-Molecular Hyperspectral PRM-SRS Imaging"

**Supplementary Figures**

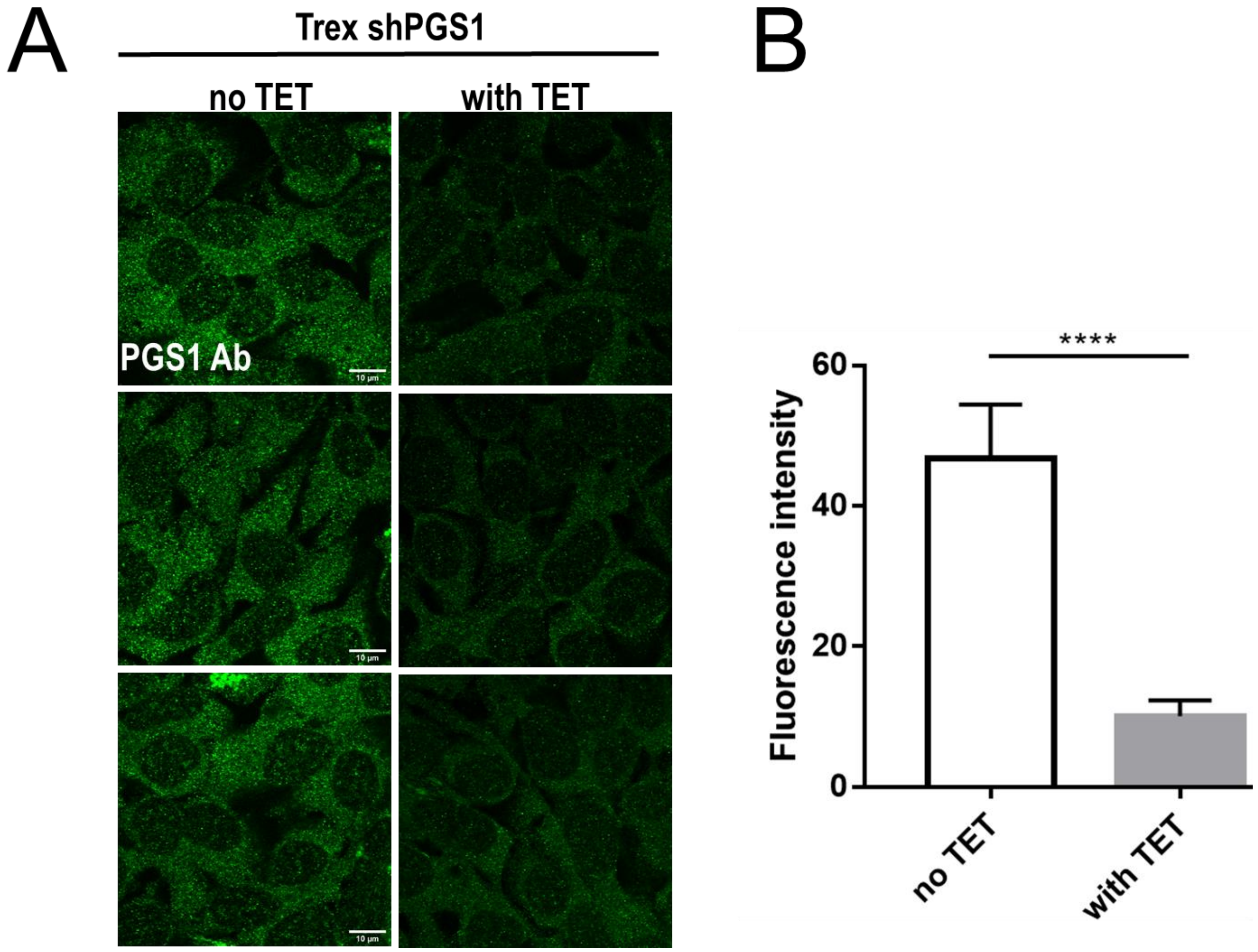


Figure S1: Immunofluorescence microscopy confirmed reduced PGS1 protein expression in shPGS1 cells. (A) Immunofluorescence staining of stable shPGS1 cells using the specific PGS1 antibody. (B) Quantification of immunofluorescence signal intensity in images shown in panel A. Compared to the control group, the PGS1 protein level was were significantly reduced by shPGS1 following induction with tetracycline (Tet).

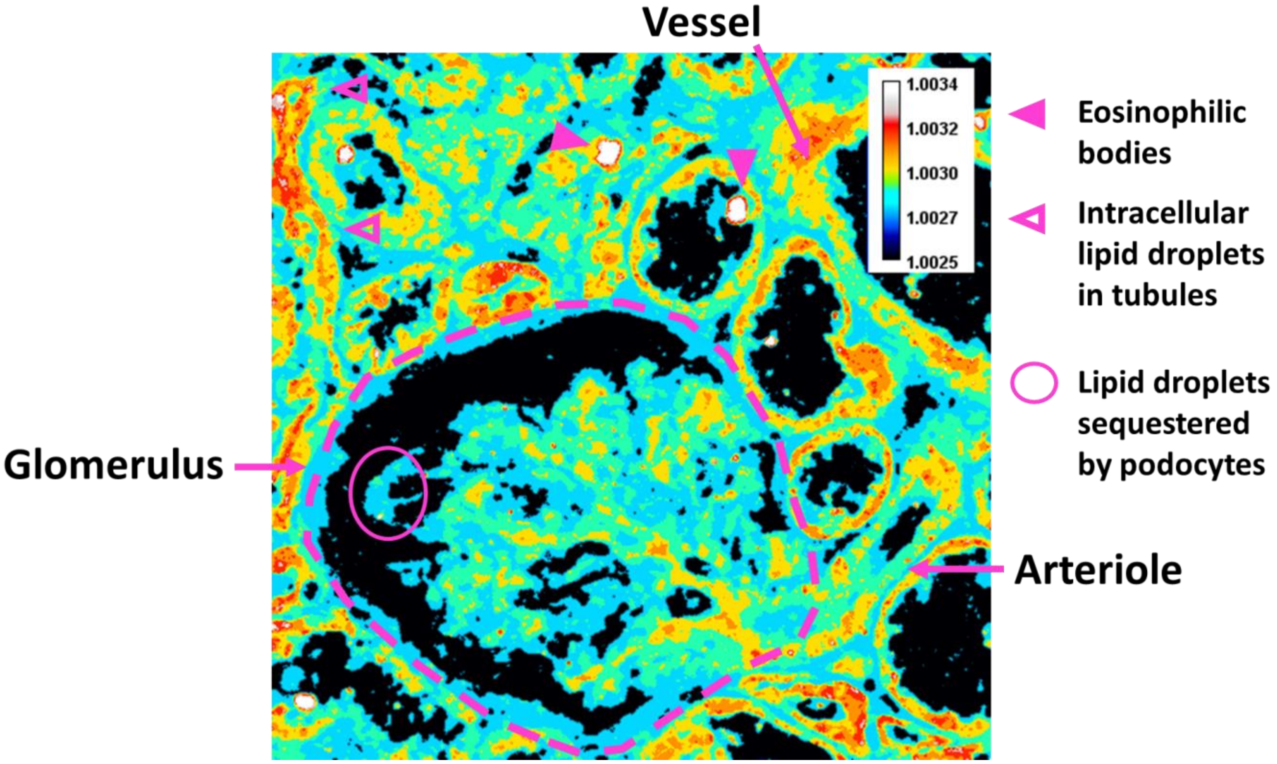


**Figure S2: Ratiometric image of cholesterol to Esterified Cholesterol corresponding to the PRM-SRS image in Figure 5B.** Ratio values are based on the similarity score ratios from the PRM-SRS images, and are not calibrated to a concentration. Higher similarity scores for cholesterol/esterified cholesterol ratio are detected in tubules and glomerular epithelial cells that line the arteriole and capillaries of the mesangium, as well as the larger deposits indicated by solid arrows in Figure 5B.

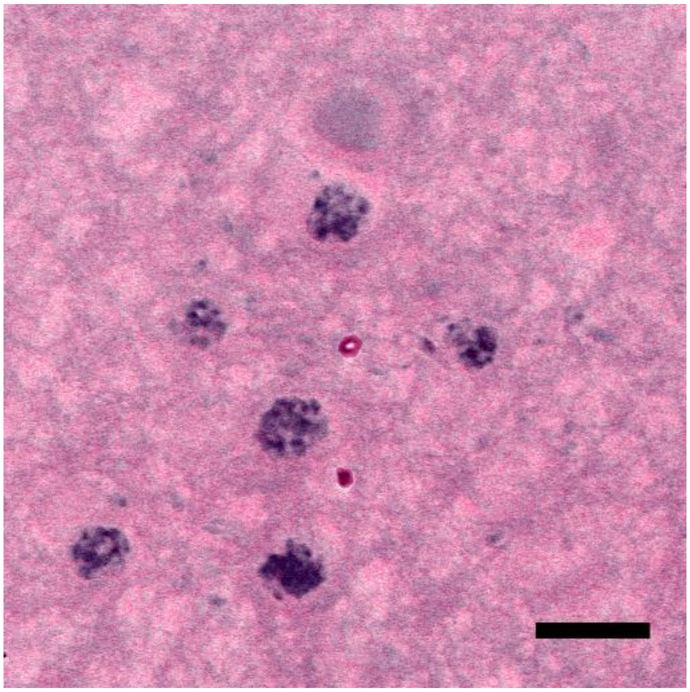

**Figure S3:** SRH (virtual H&E) image of the human brain temporal cortex sample.  Scale bar: 10 µm.

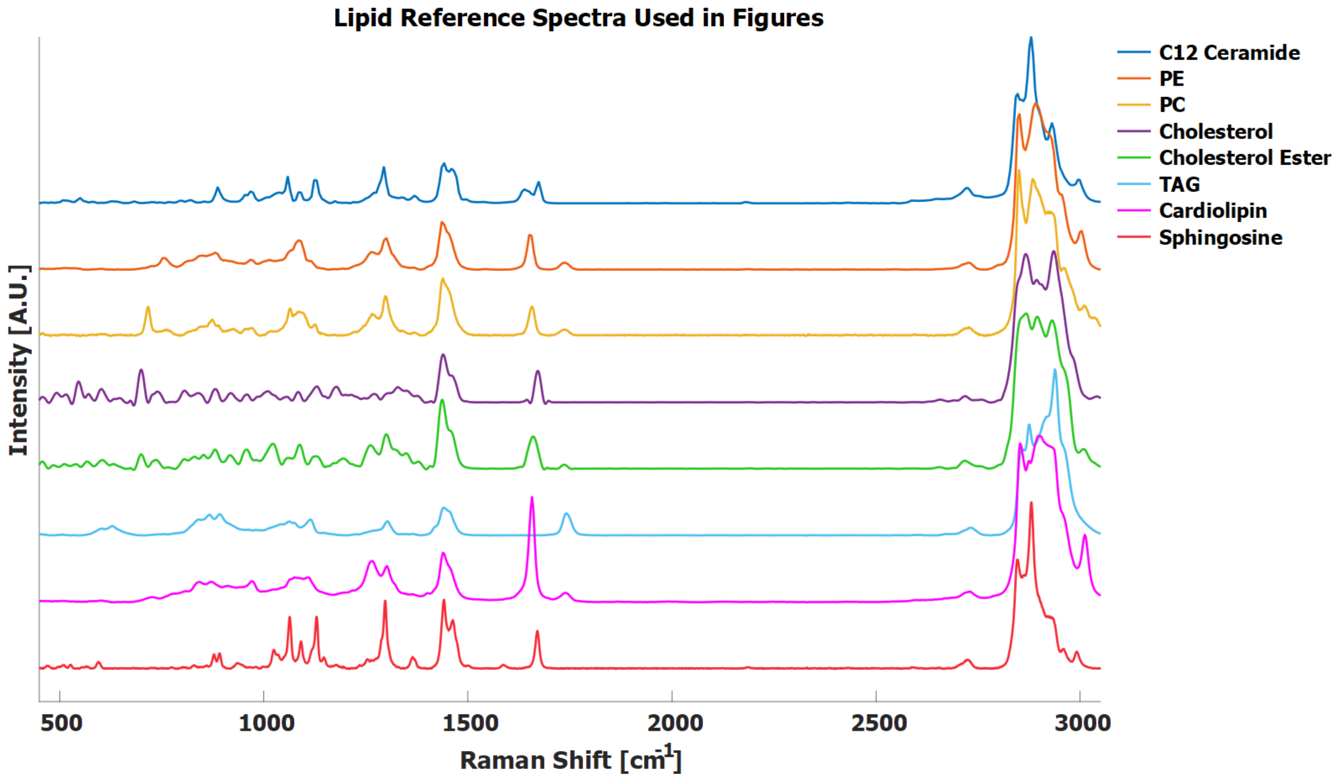


**Figure S4:** Normalized Raman spectra of different reference lipid standards used in this study.
